# ArtiaX: An Electron Tomography Toolbox for the Interactive Handling of Sub-Tomograms in UCSF ChimeraX

**DOI:** 10.1101/2022.07.26.501574

**Authors:** Utz H. Ermel, Serena M. Arghittu, Achilleas S. Frangakis

**Affiliations:** Buchmann Institute for Molecular Life Sciences and Institute for Biophysics, Goethe University Frankfurt, 60438 Frankfurt am Main, Germany; Frankfurt Institute for Advanced Studies, 60438 Frankfurt am Main, Germany

**Keywords:** cryo-electron tomography, 3D visualization, UCSF ChimeraX

## Abstract

Cryo-electron tomography analysis involves the selection of macromolecular complexes to be used for subsequent sub-tomogram averaging and structure determination. Here, we describe a plugin developed for UCSF ChimeraX that allows for the display, selection, and editing of particles within tomograms. Positions and orientations of selected particles can be manually set, modified and inspected in real-time, both on screen and in virtual reality, and exported to various file formats. The plugin allows for the parallel visualization of particles stored in several meta data lists, in the context of any 3D image that can be opened with UCSF ChimeraX. The particles are rendered in user-defined colors or using colormaps, such that individual classes or groups of particles, cross-correlation coefficients or other types of information can be highlighted to the user. The implemented functions are fast, reliable and intuitive, exploring the broad range of features in UCSF ChimeraX. They allow for a fluent human-machine interaction, which enables an effective understanding of the sub-tomogram processing pipeline, even for non-specialist users.

## Introduction

Visualization of macromolecular complexes is an inherent component of data analysis and interpretation in structural biology^1–4^. Over the last few years numerous algorithms and software packages were introduced that streamline the workflow for cryo-electron tomography (cryoET) and improve the resolution of electron density maps obtained using sub-tomogram averaging (STA)^5–10^. In addition, visualization tools were developed that provide an informative means of displaying the obtained average densities in their native context^10–13^. At the same time, advances in sample preparation techniques, such as cryogenic focused ion beam milling (cryo-FIB-SEM)^14–16^, have allowed researchers to routinely acquire tomograms of native – thicker– specimens of eukaryotic cells. Together, these developments open opportunities to study cellular architecture in unprecedented detail.

An aspect that remains challenging, in fact with all 3D images independently of the application, is the human-machine interaction during particle selection and inspection. Difficulties arise due to the 3D nature of the data, the low signal-to-noise ratio (SNR), and most significantly the missing wedge effect caused by the limited-angle tomography setting of cryoET experiments. Particle selection can therefore be a task strongly dependent on the experience and background knowledge of an expert user, especially in the case of *in situ* data. It often requires the incorporation of prior information on the particle positions, given the complex geometry of cellular structures. The selection task is addressed in several software packages, such as Dynamo^10,17^, IMOD/PEET^6,12,18^, EMAN2^9,19^, Particle Picker for UCSF Chimera^20^, the EM Toolbox^11^, and UCSF ChimeraX itself^21^, which allow manual particle picking, optionally guided by geometrical models. Other commonly employed methods for particle selection are purely computational, such as template matching (offered in Dynamo^17^, Warp/M^7,22^, pyTom^5^, emClarity^8^, EMAN2^9^ or molmatch^23,24^), or neural networks (offered by EMAN2^19^ or DeepFinder^25^). Nevertheless, they often require manual post-processing or further computational curation by 3D image classification.

Here, we present an open-source toolbox, ArtiaX, that was developed on the platform of UCSF ChimeraX (“ChimeraX” hereafter)^21,26^. Our plugin allows for a fluent human-machine interaction as well as an improved visualization of cryoET-related tasks and explores the VR capabilities of ChimeraX. The toolbox improves current software solutions in three ways: (1) At the stage of selection and inspection of putative positions of various particles for further processing—they can be selected in 2D and 3D and their orientation can be set in an efficient manner using virtual reality (VR) capabilities. (2) At the stage of inspection of particles after processing— outliers can be interactively rejected. (3) In terms of user experience—interactive real-time visualization of cryoET data is made available even to non-specialist users and without requiring specialized or commercial software, enabling sharing of data with a broader audience. Additionally, users benefit from the multitude of features of ChimeraX, such as cross-platform operation, animation capabilities, and volume file format compatibility^21,26^.

## Results

One of the stated design goals of ChimeraX is enabling straightforward development of plugins for the platform by providing a software bundle interface and tool repository^21^. ArtiaX was developed based on this interface and provides new data models, file import/export capabilities, new user interface (UI) elements (Figure 1), additional commands and new mouse/VR manipulation modes tailored to the needs of researchers in the cryoET field. The plugin allows for performant, interactive display of cryo-tomograms and associated metadata in the most common file formats (AV3/TOM package^27^ and IMOD/PEET^6^ motive lists, RELION STAR files^28,29^, and Dynamo tables^10,17^). Particle lists can be visualized, edited and stored in any available format.

**Figure 1:**
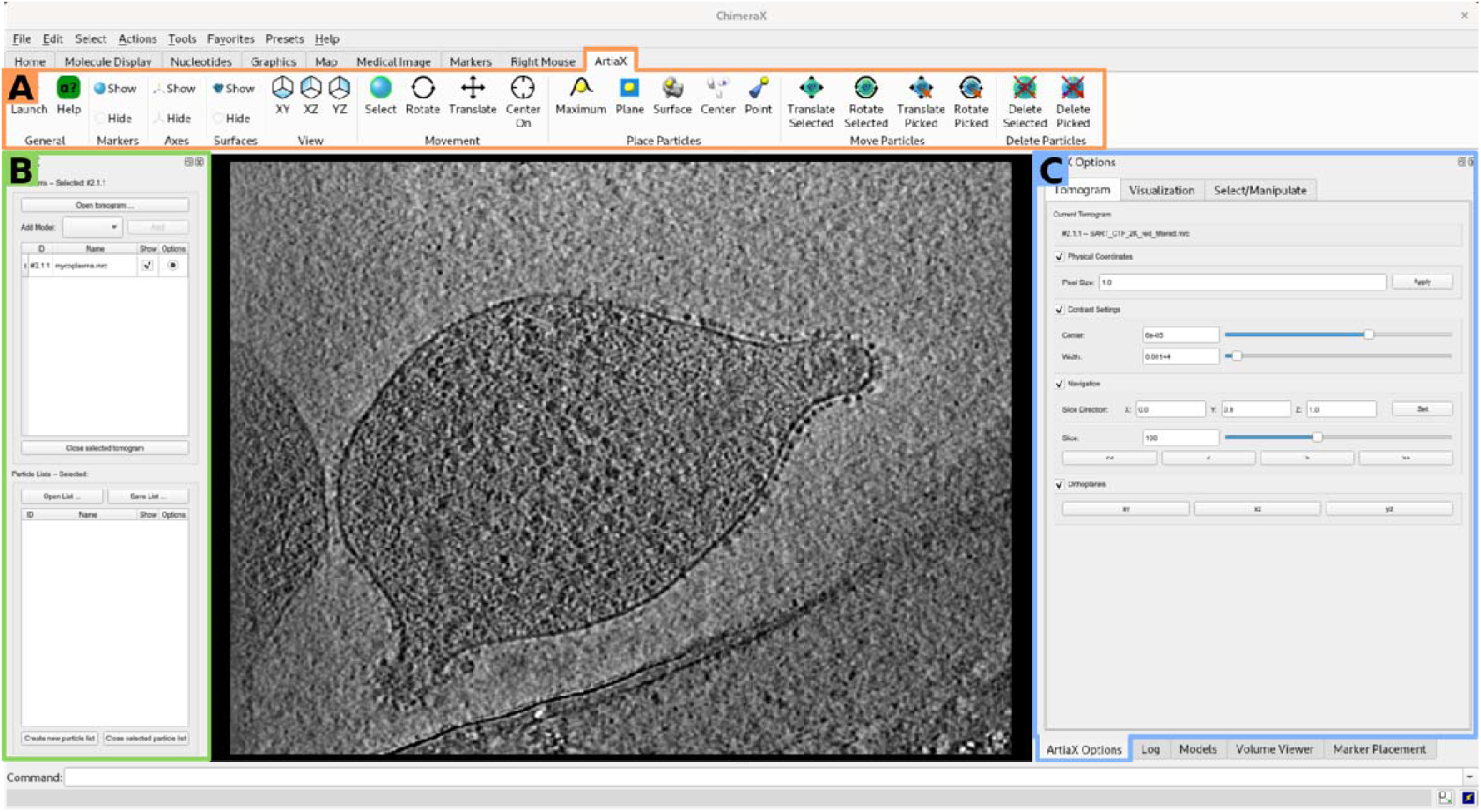
The user interface of ArtiaX within ChimeraX. (**A**) The ArtiaX toolbar provides shortcut access to frequently required operations and custom mouse modes. (**B**) Opened tomograms and particle lists are organized in a separate panel, which also allows file import and export using custom file dialogs. (**C**) The “ArtiaX Options”-panel provides UI elements for the manipulation of tomograms and particle lists (see Figure 2E-G).

### Tomogram visualization

ChimeraX supports import and rendering of electron tomograms or segmentation masks (usually in MRC-, EM- or HDF-format) in a global geometry frame based on the physical pixel size and positions stored in the file headers^21,26^. The ArtiaX plugin makes use of the platform’s volume import functionality, but maintains an internal list of tomographic volumes in order to allow easy data organization by clearly separating the tomogram models from other types of imported data. Imported tomograms are organized in a separate UI panel, in addition to the standard ChimeraX model panel (Figure 1B). Volumes can either be directly added to the plugin’s internal list of tomograms or pre-processed using built-in ChimeraX commands before import. This allows the user to make use of image processing functionality built into the platform.

Compared to ChimeraX, the ArtiaX tomogram interface provides additional UI elements enabling a number of commonly required operations. Standard viewing directions (xy-, xz- and yz-planes) can be easily recovered using shortcut buttons in the toolbar (Figure 1A). Simple navigation through tomogram planes in oblique and orthogonal directions is enabled using keyboard shortcuts and graphical sliders, allowing more precise adjustments than possible with standard mouse modes, but avoiding the need for commands (Figure 2E). Additionally, a simplified contrast control interface is provided, under the assumption of normally distributed image intensities, as they are often encountered in cryoET data.

**Figure 2:**
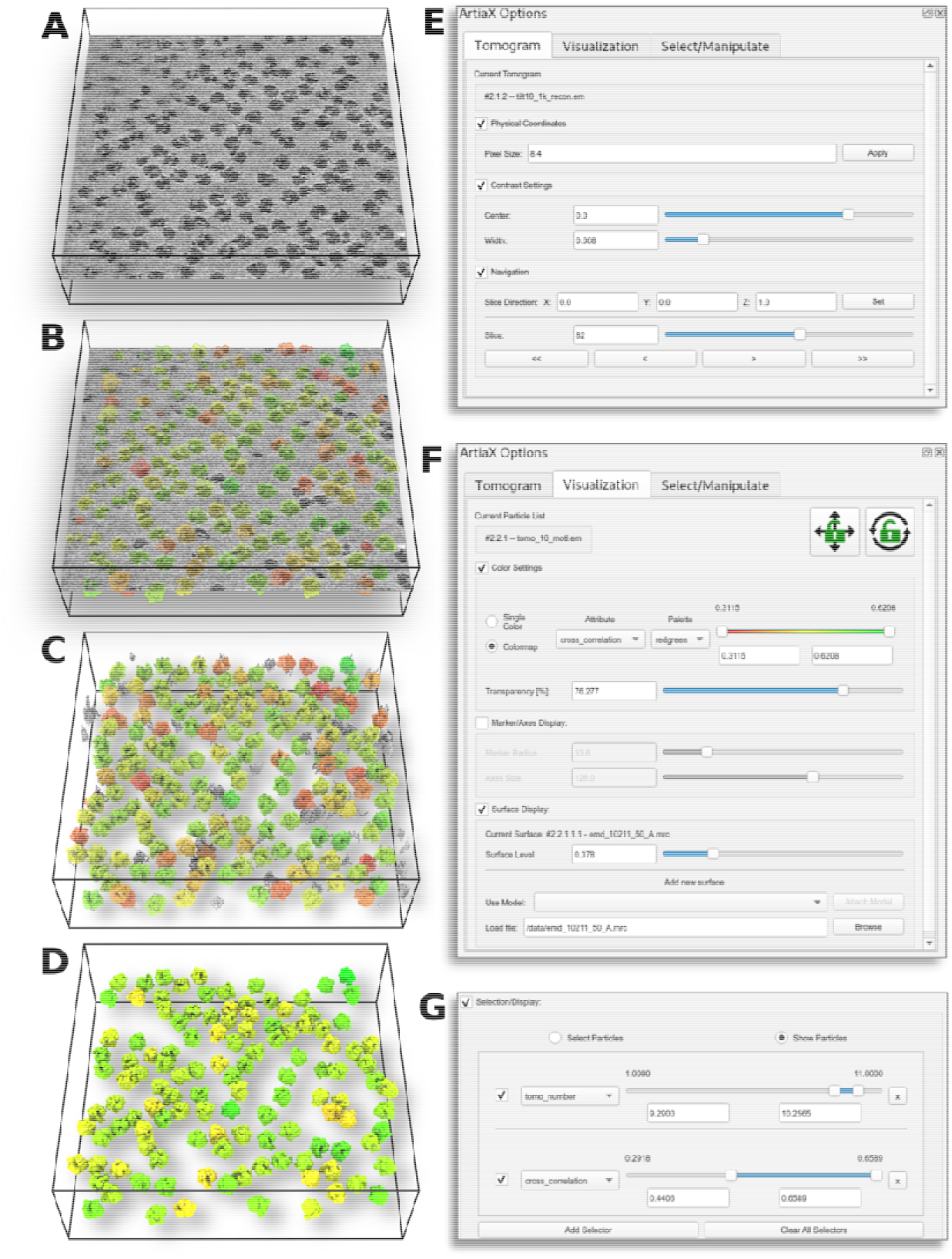
Tomogram display and metadata-based rendering. Tomographic volumes (EMPIAR-10304) are displayed as non-transparent slices by default (**A**), and contrast settings, slice direction and position are accessible in the tomogram options panel (**E**). Locations of matched *E. coli* ribosomes (EMD-10211) can be indicated transparently on either tomographic planes (**B**) or isosurface views of the tomogram (**C**). The particle visualization panel (**F**) allows adjusting the particles’ isosurface level (Surface Display box) and to set colormaps based on metadata, e.g. cross-correlation (Color Settings box). At user’s choice, rendering is limited to those particles (**D**) inside selected metadata ranges (Selection/Display box), for example tomogram number and cross-correlation (**G**).

### Particle list import and export

At present, ChimeraX already offers support for indicating image features, and links between them, via the “MarkerSet”-interface and the CMM-file format. This functionality was specifically intended for object selection in 3D images^21^ and is integrated into several cryoET processing pipelines for the purpose of particle selection^5,30^. However, the “MarkerSet”-interface models individual particles as points and is as such not designed to include or render information about the orientation of particles at each contained location. ArtiaX extends the “MarkerSet”-interface to a “ParticleList”-interface, which additionally stores and renders orientations, alignment shifts and other metadata and adds support for metadata formats other than CMM.

In the context of cryoET processing pipelines, the information on individual particles is typically stored in text-based or binary metadata files (called particle or motive-lists), which are input to or output of sub-tomogram averaging or classification procedures^5,6,8,9^. Several metadata are stored, including the 3D location of the particles within the original tomogram, alignment shifts for extracted particle images, orientations (e.g. Euler rotation angles), cross-correlation coefficients to a reference density, and many others (a detailed description of supported metadata formats can be found in the ArtiaX help files). Conventions concerning the representation of particle orientations, order of transformations and available metadata differ between file formats. Upon import into ArtiaX, particle positions, alignment shifts and orientations are converted to a common geometry convention and used to compute a transformation, which moves a reference map into the tomogram coordinate frame. Particle lists can then be exported from the common “ParticleList” structure to any of the available file formats. Like imported tomograms, “ParticleList”-models are organized in a separate panel (Figure 1B) to allow fast access to related user interface elements (Figure 2F-G).

Importantly, being an extension of the “MarkerSet”-interface, imported particle lists also maintain a regular “MarkerSet” representation of the data and are compatible with built-in marker tools and mouse modes (Figure 1A). Users can interact with “ParticleList”-models in the same ways as they did with “MarkerSet”-models, but gain the ability to set and inspect particle orientations.

### Particle list visualization

After import of particle list files, the contained particles are rendered in the 3D scene as small spheres centered at the particle location, as well as arrows indicating the particles’ orientation (Figure 3A/B). In contrast to plane-based manual picking tools (e.g. EMAN2, pyTom, or Dynamo), particles are always rendered in 3D, and can be displayed in VR immediately upon import.

**Figure 3:**
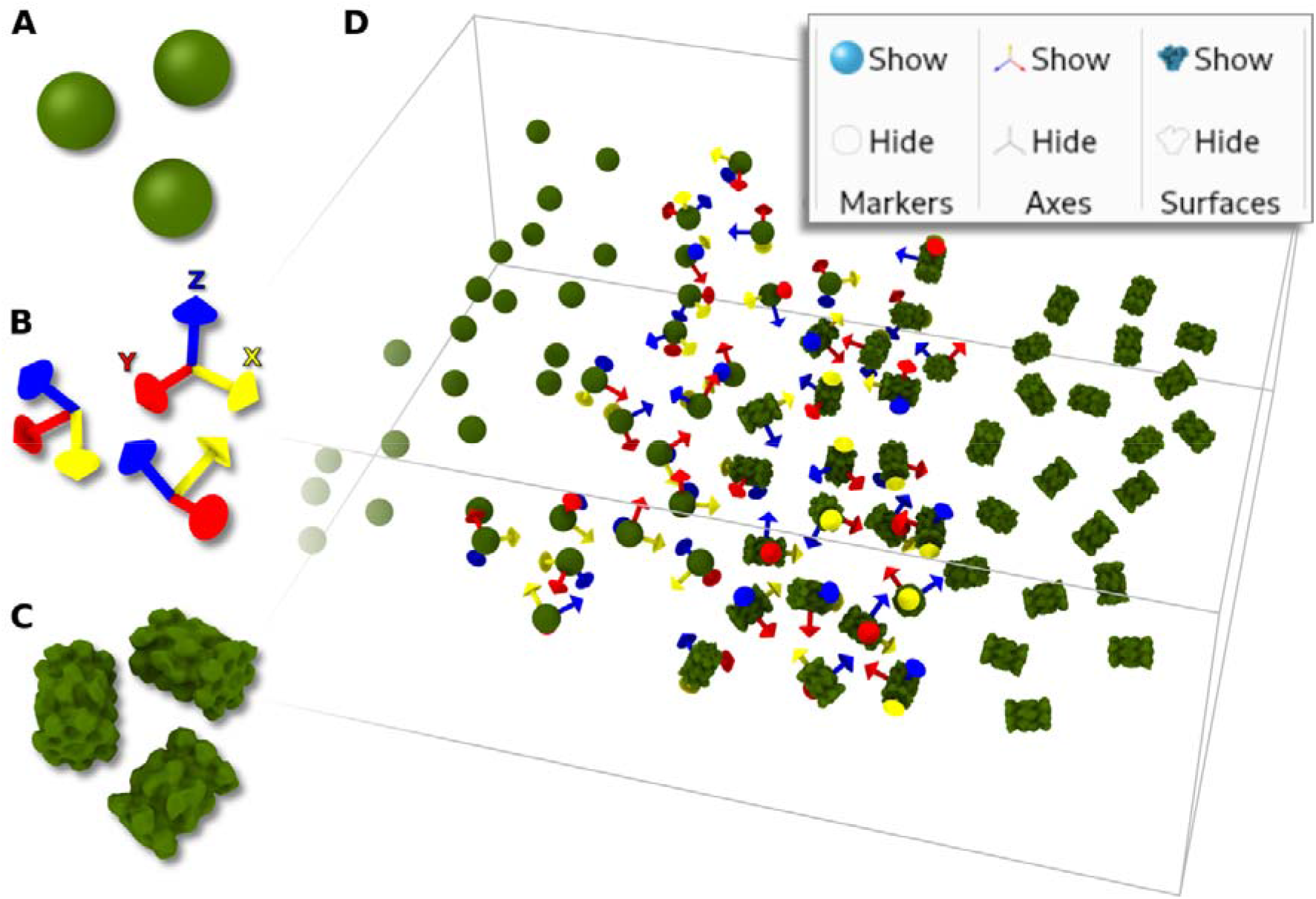
Particle list display styles. ArtiaX renders particles either as ChimeraX “marker” (**A**), using arrows indicating the particle orientation (**B**), or surfaces extracted from an associated density map (**C**), EMD-4149. Within a tomographic volume, particle locations can be rendered using any combination of these display styles (**D**) and set using display style shortcuts from the ArtiaX toolbar (top right overlay).

Metadata for each particle are stored as “attributes” in the “MarkerSet”-representation of the “ParticleList”-models. This allows selecting or hiding of subsets of the particles via built-in ChimeraX commands. Additionally, ArtiaX provides a particle selection UI, which enables interactive sub-setting based on user-defined thresholds on metadata values (Figure 2D/G). Thresholds on multiple values can easily be combined for fast and fine-grained selection, permitting, for example, to filter displayed particles for a unique tomogram ID or class, even if particles of multiple tomograms and classes are stored in the same list. Similarly, the particle list UI allows coloring particles based on metadata with user-defined colormaps (Figure 2B-D/F), which can be interactively adjusted. Combined, these features enable easy identification and removal of outliers and aid in the manual inspection of template matching, STA or classification results, within the context of tomograms (Figure 2A-D, Supplementary Video 1).

### Rendering of particle surfaces

The visualization of isosurfaces of high-resolution density maps associated with particles in the context of lower resolved tomograms is a non-trivial task. Previously, this was solved by creating intermediate, static surface representations of the particle^11^, resampling the rotated and shifted maps on the grid of the tomograms^19^, or rendering each surface individually (default for density maps in ChimeraX)^21^. While the first approach allows fast rendering, it precludes changing the isosurface level after initial surface export. The second approach leads to a loss of resolution, if the resampling grid has a lower sampling step than the reference map, or requires large amounts of storage space if the resampling grid is very fine. Lastly, while in principle display of many (>1000) copies of the same density map is possible in ChimeraX already, this incurs high computational overhead. Rendering performance measurements using a 3D scene comprising three segmented surfaces and 1720 individual density map isosurfaces of four particle species (Figure 4), indicate that performance is limited by memory throughput and CPU power (Figure 5). On four assessed hardware setups (Supplementary Table 1), rendering frame rates were measured to be below 10 frames/s, independent of the graphics processing unit (GPU), limiting real-time interactivity with the scene.

**Figure 4:**
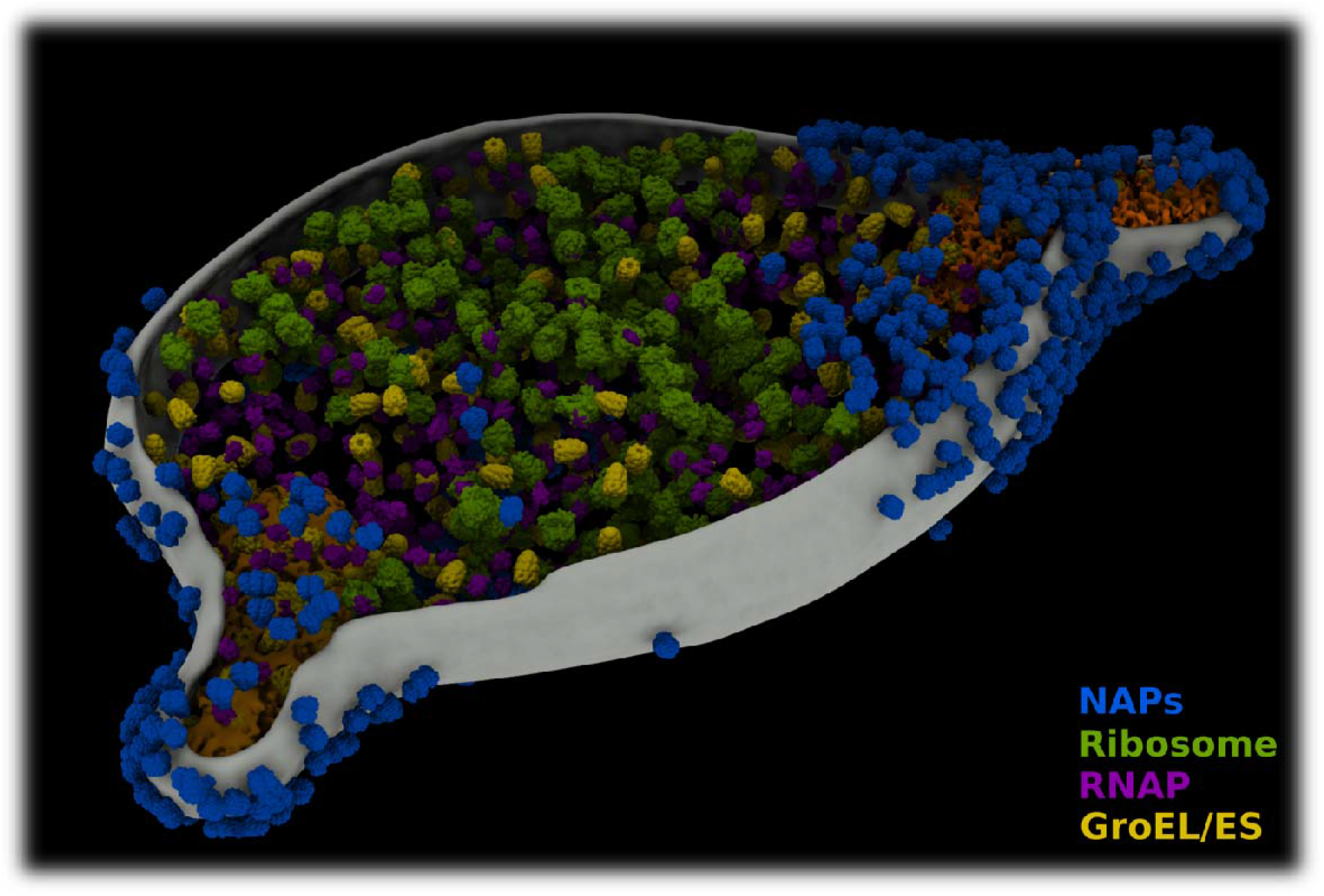
Visualization of segmentation maps and multiple particle species. Complex 3D scenes can be rendered and interactively visualized using ArtiaX. In this case, the segmented cell membrane and two terminal organelles of an *M. genitalium* cell, as well as 1720 experimental (NAPs, blue) and simulated (ribosomes, green; RNA-polymerases, purple; GroEL/ES, yellow) particles are displayed.

**Figure 5:**
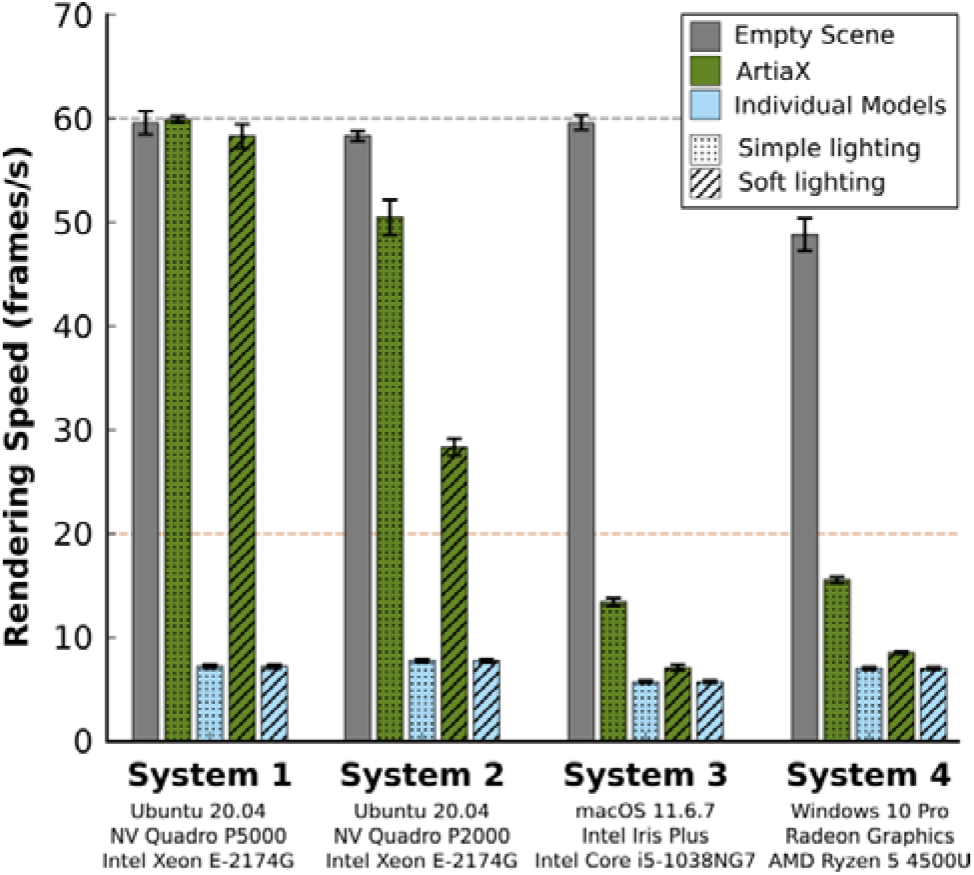
Rendering performance for complex 3D scenes. Rendering performance of the scene shown in Figure 4 is increased between 2- and 9-fold in “simple lighting”-mode (dotted bars) when using ArtiaX (green bars), compared to rendering 1720 particle surfaces as individual models (light blue bars), approaching the built-in limit (60 frames/s, grey dashed line) on high-end hardware (System 1). Visually smooth rendering (>20 frames/s, orange dashed line) is maintained in “soft-lighting”-mode (hatched bars) even on medium-range hardware (System 2). Error bars indicate standard deviation.

ArtiaX remedies this issue by introducing a novel type of model, the “SurfaceCollectionModel”, optimized for rendering many copies of the same surface in individual orientations, with high performance. As the surfaces of all particles in one “ParticleList” are assumed identical, we make use of more efficient OpenGL rendering routines, which are already employed in ChimeraX for rendering atomic structures efficiently^21^. Our implementation allows for 2- to 9-fold higher rendering frame rates when using simplified graphics settings (“simple lighting”) on all tested hardware setups (Figure 5). On medium-range to high-end hardware, visually smooth rendering (>20 frames/s) is possible using more demanding display settings (“soft lighting”). This performance improvement translates directly to VR, enabling VR visualization of large sets of particle surfaces.

At user’s choice, it is possible to place experimentally obtained density maps of any ChimeraX-supported volume format, or density maps generated from PDB files using the “molmap”-command at the location of each particle (Figure 3C-D, Figure 4, Supplementary Video 2). In contrast to previous methods using static surfaces, isosurface thresholds for map visualization can be set by the user in real-time for all locations at once. As a result, interactive visualization of multiple particle lists comprising 1000 or more complex surfaces is possible even when using consumer grade hardware or working in VR (Supplementary Video 3).

### Particle list editing

As in the case of the “MarkerSet”-interface, new particles within the tomograms can be selected either on individual tomographic planes, which are orthogonally or obliquely oriented to the principal axes of the 3D image (Figure 6A-B), or on the isosurface visualization of the data. This is particularly helpful in z, where the point spread function of the reconstruction is elongated due to the missing wedge. Typically, the particle orientations are estimated using some geometrical model based on prior knowledge about the specimen. Using new mouse modes implemented by ArtiaX, each of the visualized particles can be rotated and translated individually or in groups, with the new positions being stored and available for export (Figure 6C, Supplementary Video 4). Importantly, particle picking and editing is also possible in the ChimeraX VR mode (Supplementary Video 3). In this way two goals can be reached: (1) the pre-orientations can be precisely set and exported and (2) changes through sub-tomogram averaging can be tracked and interpreted in the context of the original tomograms.

**Figure 6:**
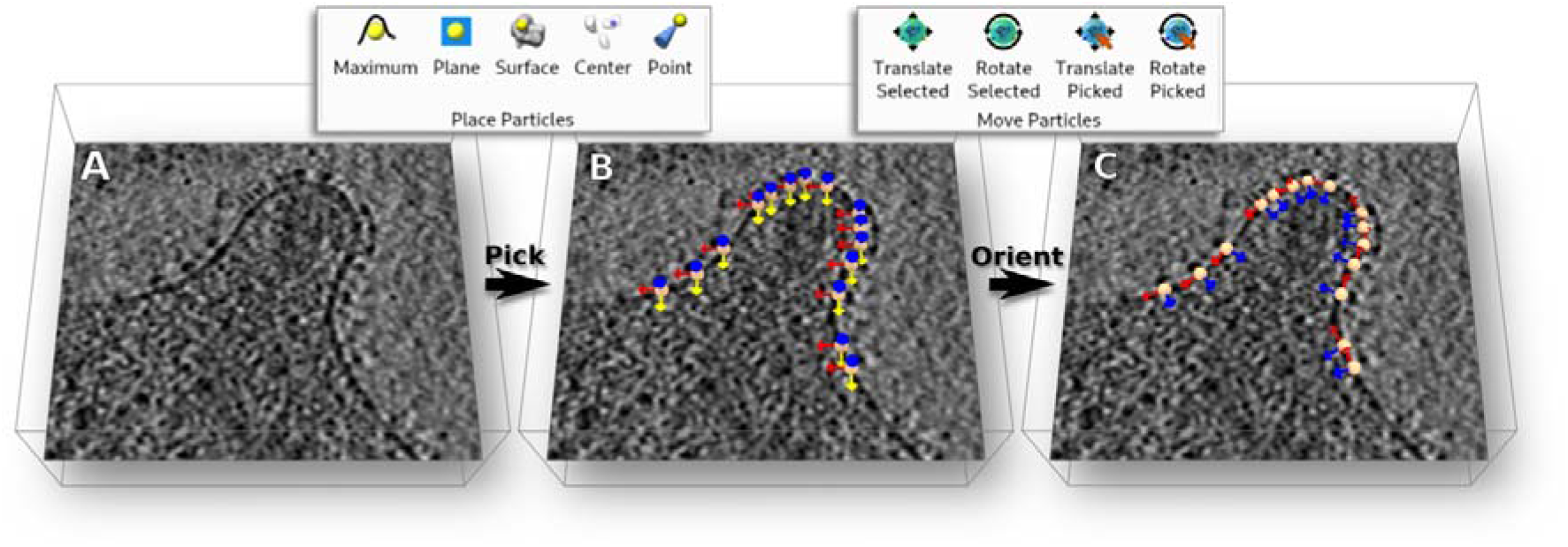
Particle picking and orientation: Individual major surface adhesion complexes can be picked on planes of a tomogram of an *M. genitalium* cell^39^ (**A**) using built-in ChimeraX mouse modes (top left overlay). Particles are initially created in default orientations at selected positions (**B**). Using the orientation mouse modes provided by ArtiaX (top right overlay), particles are oriented along contextual information inside the tomogram, in this case the cell membrane (**C**).

### Scripting capabilities

In order to facilitate recreation and sharing of 3D scenes of tomographic data created using ArtiaX, any action performed using the plugin UI can be reproduced either using built-in ChimeraX commands or newly implemented commands with simplified syntax. Additionally, ArtiaX integrates with the native file import and export commands provided by ChimeraX. Complex models involving multiple particle species (Figure 4) can be restored easily and distributed in compact fashion (Supplementary Data 1).

## Discussion

ChimeraX is a popular tool for visualization of multiscale data and associated molecular models, its utilization being widely spread throughout the structural biology community ^21,26^. Since the inception of its predecessor, UCSF Chimera^31^, the scope of both platforms has continuously expanded towards electron microscopy image data. Our plugin now adds a collection of basic capabilities to extract information out of 3D image data sets, taking into account the special needs of cryoET. This goal is achieved while maintaining a responsive user experience easily accessible to everybody, and compatibility to built-in ChimeraX functionality.

Current processing pipelines commonly store metadata in custom file formats and provide individual solutions for editing and display, sometimes limited to specific operating systems^5^. This forces researchers to rely on custom conversion solutions or familiarize themselves with a multitude of different tools providing essentially similar functionality, simply in order to assess and compare results. By providing a plugin for the already widely used, cross-platform tool ChimeraX, we hope to create a more general means for the visualization of such data.

ArtiaX does not rely on commercial platforms, such as Amira-Avizo (EM Toolbox^11^) or MATLAB (Dynamo^10^, emClarity^8^, PEET^6^). Additionally, available packages use a variety of conventions for parametrizing particle locations and orientations. Consequently, while visualization of cryoET data and manual interaction with it is an integral part of any cryoET processing pipeline, current solutions remain relatively insular and do not yet make use of the full potential of modern visualization tools, such as virtual reality.

Apart from providing a more general software solution to the cryoET field, the basic features of ArtiaX can also be used in other contexts. 3D single molecule fluorescence microscopy techniques utilize analysis methods similar to those in cryoET^32–34^ and thus require similar tools. As ChimeraX is already designed to handle fluorescence microscopy data, our plugin could be of use to the SMLM community as well, and present a contact point for integration of both methods.

Currently, our toolbox offers only a basic set of tools. While presenting immediate benefit to researchers, they concern merely one of the challenges in the cryoET field among many. Direct integration of prior knowledge in the form of easy-to-use geometric or mesh-based models in a VR context is one of our future goals. Other areas of current development include the manual and automatic segmentation of tomograms^35^, tomogram denoising^7^ and pattern recognition within tomograms^7^. In future releases, we plan to further extend ChimeraX functionality to include tools addressing these challenges. By making use of an open-source, extensible platform we also hope to invite contributions from other researchers in the field.

## Materials and Methods

### Implementation

ArtiaX was implemented in Python 3.9 according to the ChimeraX bundle development guidelines. It is packaged and distributed as a Python wheel (Supplementary Software 1). The plugin was tested on ChimeraX 1.3 and 1.4. The implementation is compatible with both Qt5 and Qt6, and as such depends on either PyQt5 or PyQt6 for the graphical user interface. Other third-party dependencies include numpy (for linear algebra routines), superqt (for additional UI elements), starfile and pandas (for parsing RELION STAR files). The bundle source code, release and installation instructions are available from https://github.com/FrangakisLab/ArtiaX under GPLv3 license.

### Tomogram reconstruction and processing

Tomograms of purified *Escherichia coli* 70S ribosomes from the publicly available EMPIAR-10304 dataset^36^ were reconstructed at a pixel size of 8.4 Å using super-sampled SART^37,38^. Particle locations and orientations were determined using template matching^23^ using the density map EMD-10211 as a reference. Particles were visualized using EMD-10211, low-pass filtered to a resolution of 50 Å.

A tomogram of a *Mycoplasma genitalium* G37 wild-type cell was reconstructed from previously published tilt series^39^ using super-sampled SART^37,38^ and binned to a pixel size of 13 Å. The cell membrane and terminal organelles were segmented using Amira 5.3.3 (ThermoFisherScientific, Waltham MA, USA). The segmentation of the cell membrane was post-processed using mean curvature motion smoothing^40^. Locations and orientations of the major surface adhesion complex (NAP) were used as determined by previous sub-tomogram alignment and averaging^41^, and visualized using the resulting average density map EMD-20259^42^, low-pass filtered to a resolution of 20 Å. Locations and orientations of ribosomes, GroEL/ES complexes and RNA-polymerases were simulated by random placement within the cell body. For visualization, surfaces were generated from PDB-7K00^43^, PDB-1PCQ^44^, and PDB-1I6V^45^, respectively, using the ChimeraX “molmap”-command at a resolution of 20 Å.

### Performance Measurements

Rendering performance was assessed for ChimeraX 1.3 using the ChimeraX “graphics rate”-command, modified to write measured frame rates to text files (Supplementary Software 2). On each assessed machine (Supplementary Table 1), a demo scene comprising three segmentation maps and 1720 particles (517 NAPs, 225 ribosomes, 364 GroEL/ES complexes and 614 RNA-polymerases) was generated using a ChimeraX script and ArtiaX commands (Supplementary Data 1). For performance comparison, an identical scene was recreated via a Python script, using individual models for each particle and relying exclusively on built-in ChimeraX functionality (Supplementary Software 2). Frame rates were measured at 1 s intervals for 100 s using “simple” and “soft” lighting settings. Baseline performance was measured using the same method with an empty scene.

## Supporting information

Supplementary Software 1

Supplementary Software 2

## Supplementary material

**Supplementary Table 1 (supplementary_table_1.docx)** – Hardware specifications of the computer systems used for measuring rendering performance.

**Supplementary Video 1 (supplementary_video_1.mp4)** – A video demonstration of particle list inspection tasks performed using ArtiaX, using a tomogram section from the EMPIAR-10304 dataset and EMD-10211 density map.

**Supplementary Video 2 (supplementary_video_2.mp4)** – A video demonstration of the assembly of a complex 3D scene using ArtiaX, comprising segmentations of an *M. genitalium* cell membrane and terminal organelles, as well as four particle lists containing 1720 particles in total.

**Supplementary Video 3 (supplementary_video_3.mp4)** – A video demonstrating particle picking, editing and removal using ArtiaX in virtual reality, as well as VR interaction with a complex 3D scene.

**Supplementary Video 4 (supplementary_video_4.mp4)** – A video demonstration of particle selection and orientation tasks performed using ArtiaX, using a tomogram of an *M. genitalium* cell.

**Supplementary Software 1 (supplementary_software_1.zip)** – The ArtiaX python wheel package for installation.

**Supplementary Software 2 (supplementary_software_2.zip)** – Python scripts used for measuring the rendering performance.

**Supplementary Data 1 (supplementary_data_1.zip)** – Demo dataset and scripts used to generate Figure 4.

## Acknowledgements

This work was supported by funding through FR 1653/14-1, FR 1653/6-3 and GRK 2566/1. The plugin was developed based on UCSF ChimeraX, developed by the Resource for Biocomputing, Visualization, and Informatics at the University of California, San Francisco, with support from National Institutes of Health R01-GM129325 and the Office of Cyber Infrastructure and Computational Biology, National Institute of Allergy and Infectious Diseases. We thank Mirko Plösser and Conny Sick for supporting software development, and Nisreen Ghanem, Alexandra N. Birtasu, Lisa Frank, and Gunnar Arctaedius for early testing and suggestions regarding user experience.

## Notes

https://github.com/FrangakisLab/ArtiaX

### Competing Interest Statement

The authors have declared no competing interest.

## References

1. Frangakis AS, Förster F (2004) Computational exploration of structural information from cryo-electron tomograms. Curr Opin Struct Biol 14:325–331.

2. Baumeister W (2002) Electron tomography: towards visualizing the molecular organization of the cytoplasm. Curr Opin Struct Biol 12:679–684.

3. Steven AC, Baumeister W (2008) The future is hybrid. J Struct Biol 163:186–195.

4. Subramaniam S, Bartesaghi A, Liu J, Bennett AE, Sougrat R (2007) Electron tomography of viruses. Curr Opin Struct Biol 17:596.

5. Hrabe T, Chen Y, Pfeffer S, Kuhn Cuellar L, Mangold AV, Förster F (2012) PyTom: A python-based toolbox for localization of macromolecules in cryo-electron tomograms and subtomogram analysis. Journal of Structural Biology 178:177–188.

6. Nicastro D, Schwartz C, Pierson J, Gaudette R, Porter ME, McIntosh JR (2006) The molecular architecture of axonemes revealed by cryoelectron tomography. Science (1979) 313:944–948.

7. Tegunov D, Xue L, Dienemann C, Cramer P, Mahamid J (2021) Multi-particle cryo-EM refinement with M visualizes ribosome-antibiotic complex at 3.5□Å in cells. Nature Methods 2021 18:218:186–193.

8. Himes BA, Zhang P (2018) emClarity: software for high-resolution cryo-electron tomography and subtomogram averaging. Nature Methods 2018 15:1115:955–961.

9. Chen M, Bell JM, Shi X, Sun SY, Wang Z, Ludtke SJ (2019) A complete data processing workflow for cryo-ET and subtomogram averaging. Nature Methods 2019 16:1116:1161–1168.

10. Castaño-Díez D, Kudryashev M, Arheit M, Stahlberg H (2012) Dynamo: A flexible, user-friendly development tool for subtomogram averaging of cryo-EM data in high-performance computing environments. Journal of Structural Biology 178:139–151.

11. Pruggnaller S, Mayr M, Frangakis AS (2008) A visualization and segmentation toolbox for electron microscopy. Journal of Structural Biology 164:161–165.

12. Kremer JR, Mastronarde DN, McIntosh JR (1996) Computer Visualization of Three-Dimensional Image Data Using IMOD. Journal of Structural Biology 116:71–76.

13. Stalling D, Westerhoff M, Hege HC (2005) amira: A Highly Interactive System for Visual Data Analysis. Visualization Handbook:749–767.

14. Hsieh C, Schmelzer T, Kishchenko G, Wagenknecht T, Marko M (2014) Practical workflow for cryo focused-ion-beam milling of tissues and cells for cryo-TEM tomography. Journal of Structural Biology 185:32–41.

15. Marko M, Hsieh C, Schalek R, Frank J, Mannella C (2007) Focused-ion-beam thinning of frozen-hydrated biological specimens for cryo-electron microscopy. Nature Methods 2007 4:34:215–217.

16. Rigort A, Bäuerlein FJB, Villa E, Eibauer M, Laugks T, Baumeister W, Plitzko JM (2012) Focused ion beam micromachining of eukaryotic cells for cryoelectron tomography. Proc Natl Acad Sci U S A 109:4449–4454.

17. Castaño-Díez D, Kudryashev M, Stahlberg H (2017) Dynamo Catalogue: Geometrical tools and data management for particle picking in subtomogram averaging of cryo-electron tomograms. Journal of Structural Biology 197:135–144.

18. Heumann JM, Hoenger A, Mastronarde DN (2011) Clustering and variance maps for cryo-electron tomography using wedge-masked differences. Journal of Structural Biology 175:288–299.

19. Tang G, Peng L, Baldwin PR, Mann DS, Jiang W, Rees I, Ludtke SJ (2007) EMAN2: An extensible image processing suite for electron microscopy. Journal of Structural Biology 157:38–46.

20. Qu K, Ke Z, Zila V, Anders-Össwein M, Glass B, Mücksch F, Müller R, Schultz C, Müller B, Kräusslich HG, et al. (2021) Maturation of the matrix and viral membrane of HIV-1. Science (1979) 373:700–704.

21. Pettersen EF, Goddard TD, Huang CC, Meng EC, Couch GS, Croll TI, Morris JH, Ferrin TE (2021) UCSF ChimeraX: Structure visualization for researchers, educators, and developers. Protein Science 30:70–82.

22. Tegunov D, Cramer P (2019) Real-time cryo-electron microscopy data preprocessing with Warp. Nature Methods 2019 16:1116:1146–1152.

23. Frangakis AS, Böhm J, Förster F, Nickell S, Nicastro D, Typke D, Hegerl R, Baumeister W (2002) Identification of macromolecular complexes in cryoelectron tomograms of phantom cells. Proc Natl Acad Sci U S A 99:14153.

24. Förster F, Han BG, Beck M (2010) Visual Proteomics. Methods in Enzymology 483:215–243.

25. Moebel E, Martinez-Sanchez A, Lamm L, Righetto RD, Wietrzynski W, Albert S, Larivière D, Fourmentin E, Pfeffer S, Ortiz J, et al. (2021) Deep learning improves macromolecule identification in 3D cellular cryo-electron tomograms. Nature Methods 2021 18:1118:1386–1394.

26. Goddard TD, Huang CC, Meng EC, Pettersen EF, Couch GS, Morris JH, Ferrin TE (2018) UCSF ChimeraX: Meeting modern challenges in visualization and analysis. Protein Science 27:14–25.

27. Förster F, Medalia O, Zauberman N, Baumeister W, Fass D (2005) Retrovirus envelope protein complex structure in situ studied by cryo-electron tomography. Proc Natl Acad Sci U S A 102:4729–4734.

28. Zivanov J, Otón J, Ke Z, Qu K, Morado D, Castaño-Díez D, Kügelgen A von, Bharat TAM, Briggs JAG, Scheres SHW (2022) A Bayesian approach to single-particle electron cryo-tomography in RELION-4.0. bioRxiv:2022.02.28.482229.

29. Bharat TAM, Russo CJ, Löwe J, Passmore LA, Scheres SHW (2015) Advances in Single-Particle Electron Cryomicroscopy Structure Determination applied to Sub-tomogram Averaging. Structure 23:1743–1753.

30. Scaramuzza S, Castaño-Díez D (2021) Step-by-step guide to efficient subtomogram averaging of virus-like particles with Dynamo. PLOS Biology 19:e3001318.

31. Pettersen EF, Goddard TD, Huang CC, Couch GS, Greenblatt DM, Meng EC, Ferrin TE (2004) UCSF Chimera—A visualization system for exploratory research and analysis. Journal of Computational Chemistry 25:1605–1612.

32. Heydarian H, Joosten M, Przybylski A, Schueder F, Jungmann R, Werkhoven B van, Keller-Findeisen J, Ries J, Stallinga S, Bates M, et al. (2021) 3D particle averaging and detection of macromolecular symmetry in localization microscopy. Nature Communications 2021 12:112:1–9.

33. Sabinina VJ, Hossain MJ, Hériché JK, Hoess P, Nijmeijer B, Mosalaganti S, Kueblbeck M, Callegari A, Szymborska A, Beck M, et al. (2021) Three-dimensional superresolution fluorescence microscopy maps the variable molecular architecture of the nuclear pore complex. Molecular Biology of the Cell 32:1523–1533.

34. Broeken J, Johnson H, Lidke DS, Liu S, Nieuwenhuizen RPJ, Stallinga S, Lidke KA, Rieger B (2015) Resolution improvement by 3D particle averaging in localization microscopy. Methods and Applications in Fluorescence 3:014003.

35. Chen M, Dai W, Sun SY, Jonasch D, He CY, Schmid MF, Chiu W, Ludtke SJ (2017) Convolutional neural networks for automated annotation of cellular cryo-electron tomograms. Nature Methods 2017 14:10 [Internet] 14:983–985. Available from: https://www.nature.com/articles/nmeth.4405

36. Eisenstein F, Danev R, Pilhofer M (2019) Improved applicability and robustness of fast cryo-electron tomography data acquisition. Journal of Structural Biology 208:107–114.

37. Kunz M, Frangakis AS (2017) Three-dimensional CTF correction improves the resolution of electron tomograms. Journal of Structural Biology 197:114–122.

38. Kunz M, Frangakis AS (2014) Super-sampling SART with ordered subsets. Journal of Structural Biology 188:107–115.

39. Seybert A, Gonzalez-Gonzalez L, Scheffer MP, Lluch-Senar M, Mariscal AM, Querol E, Matthaeus F, Piñol J, Frangakis AS (2018) Cryo-electron tomography analyses of terminal organelle mutants suggest the motility mechanism of Mycoplasma genitalium. Molecular Microbiology 108:319–329.

40. Frangakis AS (2022) Mean curvature motion facilitates the segmentation and surface visualization of electron tomograms. Journal of Structural Biology 214:107833.

41. Scheffer MP, Gonzalez-Gonzalez L, Seybert A, Ratera M, Kunz M, Valpuesta JM, Fita I, Querol E, Piñol J, Martín-Benito J, et al. (2017) Structural characterization of the NAP; the major adhesion complex of the human pathogen Mycoplasma genitalium. Molecular Microbiology 105:869–879.

42. Aparicio D, Scheffer MP, Marcos-Silva M, Vizarraga D, Sprankel L, Ratera M, Weber MS, Seybert A, Torres-Puig S, Gonzalez-Gonzalez L, et al. (2020) Structure and mechanism of the Nap adhesion complex from the human pathogen Mycoplasma genitalium. Nature Communications 2020 11:111:1–10.

43. Watson ZL, Ward FR, Méheust R, Ad O, Schepartz A, Banfield JF, Cate JHD (2020) Structure of the bacterial ribosome at 2 Å resolution. Elife 9:1–62.

44. Chaudhry C, Farr GW, Todd MJ, Rye HS, Brunger AT, Adams PD, Horwich AL, Sigler PB (2003) Role of the gamma-phosphate of ATP in triggering protein folding by GroEL-GroES: function, structure and energetics. EMBO J 22:4877–4887.

45. Campbell EA, Korzheva N, Mustaev A, Murakami K, Nair S, Goldfarb A, Darst SA (2001) Structural mechanism for rifampicin inhibition of bacterial rna polymerase. Cell 104:901–912.

