## Supplementary Table 1 for "ArtiaX: An Electron Tomography Toolbox for the Interactive Handling of Sub-Tomograms in UCSF ChimeraX"

**Supplementary Table 1:** Hardware specifications of 4 systems used for benchmarking rendering performance. MFCTR – manufacturer.

|  | MFCTR | CPU | RAM | GPU | OS |
| --- | --- | --- | --- | --- | --- |
| system 1 | Dell | Intel Xeon E-2174G | 64 GB | NVIDIA Quadro P5000 | Ubuntu 20.04 |
| system 2 | Dell | Intel Xeon E-2174G | 64 GB | NVIDIA Quadro P5000 | Ubuntu 20.04 |
| System 3 | Apple | Intel Core i5-1038NG7 | 16 GB | Intel Iris Plus | macOS 11.6.7 |
| system 4 | Lenovo | AMD Ryzen 4 4500U | 16 GB | Radeon Graphics | Windows 10 Pro |
