## Supplementary Software 1 for "ArtiaX: An Electron Tomography Toolbox for the Interactive Handling of Sub-Tomograms in UCSF ChimeraX": INSTALL

1. Download the latest Version of ChimeraX (version >= 1.3) to your operating system from here: 
https://www.rbvi.ucsf.edu/chimerax/download.html#release. 

2. Open ChimeraX and install the package using the command:
>toolshed install supplementary_software_1/ChimeraX_ArtiaX-0.1-py3-none-any.whl

3. Relaunch ChimeraX.
