## Supplementary Software 2 for "ArtiaX: An Electron Tomography Toolbox for the Interactive Handling of Sub-Tomograms in UCSF ChimeraX": copy_to_chimerax.py

```
from chimerax.core.generic3d import Generic3DModel

for i in range(0,4):
    pp = session.ArtiaX.partlists.get(i).collection_model.child_positions
    s = session.ArtiaX.partlists.get(i).display_model.get(0).surfaces[0]

    parent_model = Generic3DModel('ChimeraX {}'.format(session.ArtiaX.partlists.get(i)), session)
    session.models.add([parent_model])

    vertices = s._vertices
    normals = s._normals
    triangles = s._triangles
    edge_mask = s._edge_mask
    triangle_mask = s._triangle_mask


    for idx, p in enumerate(pp):
        standalone_surface = Generic3DModel('Child {}'.format(idx), session)
        standalone_surface.set_geometry(vertices, normals, triangles, edge_mask, triangle_mask)
        standalone_surface.position = p.copy()
        standalone_surface.color = session.ArtiaX.partlists.get(i).color
        parent_model.add([standalone_surface])

print('done')
```
