## Supplementary Software 2 for "ArtiaX: An Electron Tomography Toolbox for the Interactive Handling of Sub-Tomograms in UCSF ChimeraX": record_framerate.py

```
import numpy as np
import time
import csv
import sys
import threading
from pathlib import Path

from chimerax.core.commands import run

# Report frame rate.
class FrameRateReporter:
    def __init__(self, session, N, file):
        self.session = session
        self.report_interval = 1.0	# seconds
        self._new_frame_handler = None
        self._frame_drawn_handler = None
        self._num_frames = 0
        self._last_time = None
        self._cumulative_times = {'draw_time': 0,
                                  'new_frame_time': 0,
                                  'atomic_check_for_changes_time': 0,
                                  'drawing_change_time': 0,
                                  'clip_time': 0}

        self.period = N
        self.fps = np.zeros((N,))
        self.count = 0
        self.file = file

    def report(self, enable):
        t = self.session.triggers
        if enable:
            if self._new_frame_handler is None:
                self._new_frame_handler = t.add_handler('new frame', self.new_frame_cb)
                self._frame_drawn_handler = t.add_handler('frame drawn', self.frame_drawn_cb)
        else:
            nfh, fdh = self._new_frame_handler, self._frame_drawn_handler
            if nfh:
                t.remove_handler(nfh)
            if fdh:
                t.remove_handler(fdh)
            self._new_frame_handler = self._frame_drawn_handler = None
            
    def new_frame_cb(self, *ignore):
        # Make frames draw continuously even if scene does not change
        self.session.main_view.redraw_needed = True

    def frame_drawn_cb(self, *ignore):
        from time import time
        t = time()
        lt = self._last_time
        if lt is None:
            self._last_time = t
            self._num_frames = 0
            return
        self.record_times()
        self._num_frames += 1
        nf = self._num_frames
        dt = t - lt
        if dt > self.report_interval:
            self.count += 1

            if self.count >= self.period:
                self.report(False)
                
                with open(self.file, 'w', newline='') as csvfile:
                    writer = csv.writer(csvfile, delimiter=' ')
                    l = self.fps.tolist()[1:]
                    print(l)
                    for val in l:
                        writer.writerow([val])
                return

            fps = nf / dt
            self.fps[self.count] = fps
            #print(fps)
            #msg = '%.1f frames per second' % fps
            #ct = self._cumulative_times
            #msg += ', ' + ', '.join(['%s %.0f%%' % (k[:-5], 100*v/dt) for k,v in ct.items()])
            #self.session.logger.status(msg)
            self._last_time = t
            self._num_frames = 0
            ct = self._cumulative_times
            for k in ct.keys():
                ct[k] = 0

    def record_times(self):
        u = self.session.update_loop
        ct = self._cumulative_times
        for k in ct.keys():
            ct[k] += getattr(u, 'last_'+k)


N = int(sys.argv[1])

frp = FrameRateReporter(session, N, sys.argv[2])
frp.report(True)
```
